# Autologous iPSC- and MSC-derived Chondrocyte Implants for Cartilage Repair in a Miniature Pig Model

**DOI:** 10.1101/2023.07.25.550409

**Authors:** Ming-Song Lee, Athillesh Sivapatham, Ellen M. Leiferman, Hongli Jiao, Yan Lu, Brett W. Nemke, Matthew Leiferman, Mark D. Markel, Wan-Ju Li

## Abstract

Induced pluripotent stem cell (iPSC)-derived mesenchymal stem cells (iMSCs) have greater potential for generating chondrocytes without hypertrophic and fibrotic phenotypes compared to bone marrow-derived mesenchymal stem/stromal cells (BMSCs). However, there is a lack of research demonstrating the use of autologous iMSCs for repairing articular chondral lesions in large animal models. In this study, we aimed to evaluate the effectiveness of autologous miniature pig (minipig) iMSC-chondrocyte (iMSC-Ch)-laden implants in comparison to autologous BMSC-chondrocyte (BMSC-Ch)-laden implants for cartilage repair in porcine femoral condyles. iMSCs and BMSCs were seeded into fibrin glue/nanofiber constructs and cultured with chondrogenic induction media for 7 days before implantation. To assess the regenerative capacity of the cells, 19 skeletally mature Yucatan minipigs were randomly divided into microfracture control, acellular scaffold, iMSC, and BMSC subgroups. A cylindrical defect measuring 7 mm in diameter and 0.6 mm in depth was created on the articular cartilage surface without violating the subchondral bone. The defects were then left untreated or treated with acellular or cellular implants. Both cellular implant-treated groups exhibited enhanced joint repair compared to the microfracture and acellular control groups. Immunofluorescence analysis yielded significant findings, showing that cartilage treated with iMSC-Ch implants exhibited higher expression of COL2A1 and minimal to no expression of COL1A1 and COL10A1, in contrast to the BMSC-Ch-treated group. This indicates that the iMSC-Ch implants generated more hyaline cartilage-like tissue compared to the BMSC-Ch implants. These results contribute to filling the knowledge gap regarding the potential of autologous iPSC derivatives for cartilage repair in translational animal models.

## Introduction

Various types of stem cells have been explored for articular cartilage regeneration.^1^ For instance, bone marrow stromal cells (BMSCs) have been extensively studied for their ability to promote cartilage repair due to their chondrogenic capacity and ease of harvesting from patients. The microfracture procedure, which harnesses the regenerative capacity of endogenous BMSCs, is clinically used for cartilage lesion repair.^2^ This procedure induces tissue repair by exposing subchondral BMSCs at the joint surface, facilitating cartilage regeneration. However, the repaired tissue primarily consists of fibrocartilage with a high ratio of collagen type 1 to collagen type 2 and a low amount of proteoglycans.^3, 4^ Such tissue composition results in inferior mechanical properties for load resistance of the joint.^5^ Additionally, the use of BMSCs for articular cartilage repair raises concerns about osteochondral ossification of the regenerated tissue. Research evidence has shown that BMSC-derived chondrocytes express hypertrophic chondrocyte-associated markers, such as RUNX2 and COL10A1, and undergo the process of endochondral ossification.^6^ These findings suggest that BMSCs may not be suitable for hyaline cartilage regeneration.

Induced pluripotent stem cells (iPSCs), reprogrammed from somatic cells through the overexpression of Yamanaka factors, possess similar capabilities of self-renewal and lineage cell differentiation as embryonic stem cells.^7^ iPSCs serve as an *ex vivo* cell source without supply limitations and can be differentiated into any cell type, including chondrocytes, for therapeutic purposes without ethical concerns. One major advantage of iPSCs over adult tissue-derived chondrocytes is their extensive expansion potential without noticeable cellular senescence. Our recent findings, demonstrating that iPSC-derived mesenchymal stem/stromal cells (iMSCs) exhibit greater chondrogenic differentiation capacity and cell proliferation with attenuated p53/p21^CIP1^ activity compared to parental synovial fluid-derived MSCs,^8^ indicate that iPSCs are a cell source with great regenerative potential. Notably, iMSCs hold promise for generating naive neocartilage without a tendency for ossification.^9,10^ For example, Diederichs et al. compared the differences in chondrogenesis among iMSCs, BMSCs, and articular chondrocytes (ACs) and found that iMSC-derived chondrocytes did not undergo hypertrophic differentiation and were able to produce twofold more proteoglycans than BMSC-derived chondrocytes and control ACs.^11^ These strengths have positioned iPSCs as a promising source for chondrocyte derivation in articular cartilage regeneration.

Developing autologous iPSC-based therapies has been viewed as an appealing strategy due to safety concerns. Although the allogeneic strategy using “off-the-shelf” cells or tissues is more practical for preparing grafts for implantation, concerns remain regarding the immune response against implanted cells or tissues. Studies have shown that allogeneic cartilage was infiltrated after transplantation in mice^12^ and implanted allogeneic chondrocytes in a rabbit joint defect were destroyed due to a humoral immune response.^13^ These results indicate that allogeneic implanted chondrocytes were attacked by host T cells and natural killer cells detecting the expression of class II major histocompatibility complex molecules and natural cytotoxicity receptors, respectively, on the chondrocytes.^14,15^ Considering the safety of cell therapies, autologous stem cells should be a more viable source than allogeneic stem cells for cartilage regeneration.

The potential of iPSC derivatives, including MSCs and chondrocytes, for articular cartilage regeneration has been explored in small mammal models, such as rodents and rabbits.^16–18^ Although small animal models offer advantages in terms of cost, size, life cycle, and ease of maintenance over large animal models, the spontaneous cartilage repair observed in small animals due to their superior intrinsic healing ability may not be reproducible in studies involving large animals or clinical trials.^19,20^ On the other hand, large animal models, such as miniature pigs (minipigs), exhibit similarities in size, thickness, and anatomical structure of articular cartilage to humans.^21^ Therefore, minipigs are considered a desirable animal model for translational research on cartilage regeneration.

While allogeneic and xenogeneic iPSCs have demonstrated the ability to generate cartilage *in vitro* and *in vivo*,^22, 23^ it remains unclear whether it would be advantageous, in terms of clinical relevance, to use autologous iPSCs for deriving therapeutic chondrocytes for articular cartilage repair in a translational animal model. In this study, we evaluated the implantation of autologous iMSC-derived chondrocytes (iMSC-Chs) and gold-standard autologous BMSC-derived hondrocytes (BMSC-Chs) for cartilage repair of the femoral condyles in skeletally matured minipigs.

## Results

### Pluripotent characteristics of minipig fibroblast-derived iPSCs

To generate minipig iPSCs, we isolated fibroblasts from the ears of 6 animal and transfected them with the episomal plasmid pMaster12 to ectopically overexpress pluripotency factors for cellular reprogramming. At day 1 after transfection, the fibroblasts exhibited an elongated morphology, gradually transitioning into iPSCs with epithelial morphology. By day 16, tightly packed ESC-like colonies with well-defined borders had formed (Fig. 1A). After 21 days, we hand-picked the iPSC colonies and transferred them to culture plates coated with a feeder layer of mouse embryonic fibroblasts to maintain the established cell lines.

**Figure 1.**
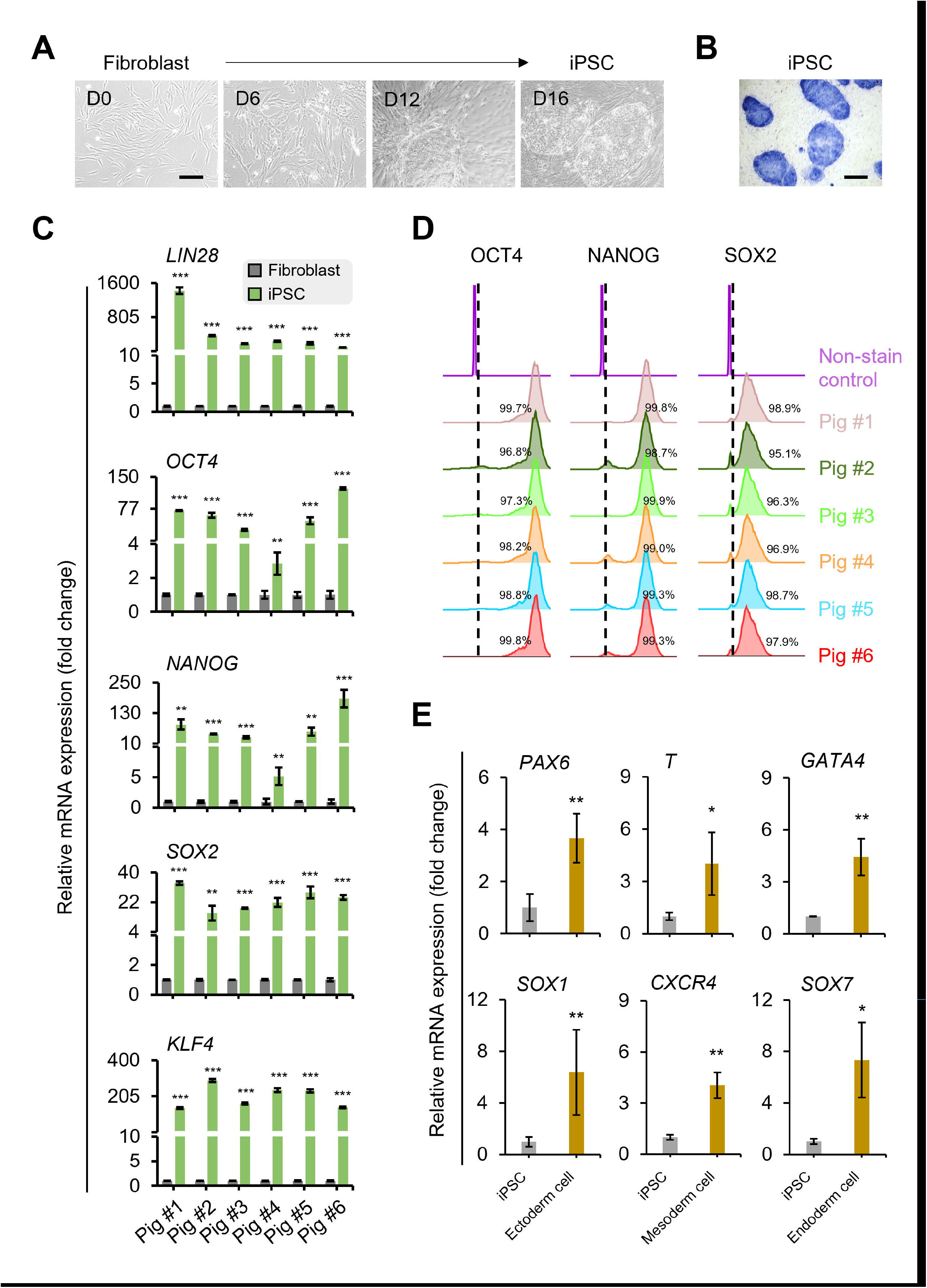
Characterization of minipig iPSCs reprogrammed from dermal fibroblasts. (A) Changes in cell morphology from minipig fibroblasts to iPSCs during cellular reprogramming between days 0 and 16. (B) Representative iPSC colonies stained with alkaline phosphatase for determination of pluripotency. (C) Transcript levels of pluripotency markers in fibroblasts and iPSCs individually derived from 6 minipigs. (D) Flow cytometry analysis of expression of pluripotency markers in 6 independent iPSC lines. (E) Transcript levels of ectodermal markers *PAX6* and *SOX1*, mesodermal markers *T* and *CXCR4*, and endodermal markers *GATA4* and *SOX7*. Scale bar = 200 µm. \**p* < 0.05, \*\**p* < 0.01, \*\*\**p* < 0.001; n = 6.

To characterize the iPSCs, we performed staining for alkaline phosphatase (ALP). The iPSC colonies consistently showed positive ALP staining (Fig. 1B). We further examined the expression levels of pluripotency markers using quantitative polymerase chain reaction (qPCR) and flow cytometry. The transcript expression of *LIN28*, *OCT4*, *NANOG*, *SOX2*, and *KLF4* was significantly upregulated in iPSCs compared to their parental fibroblasts (Fig. 1C). Flow cytometry analysis revealed that over 95% of the iPSCs derived from each minipig showed positive staining for the markers OCT4, NANOG, and SOX2 (Fig. 1D). In addition to exhibiting pluripotent characteristics, the iPSCs demonstrated the ability to differentiate into cells of the three germ layers. After 7 days of induction with lineage-specific differentiation medium, the minipig iPSCs expressed significantly higher levels of ectoderm-associated markers *PAX6* and *SOX1*, mesoderm-associated markers *T* and *CXCR4*, and endoderm-associated markers *GATA4* and *SOX7* compared to non-induced iPSC controls (Fig. 1E). Taken together, these results indicate that using our protocol, we successfully generated autologous minipig iPSCs from fibroblasts.

### Multilineage differentiation capability of iMSCs and BMSCs

Minipig iPSCs were induced to differentiate into iMSCs using the STEMdiffTM Mesenchymal Progenitor kit, while BMSCs were isolated from bone marrow following our laboratory protocol. Morphology analysis revealed that both iMSCs and BMSCs exhibited a similar spindle shape (Fig. 2A). In terms of cell growth, iMSCs displayed a higher proliferation rate in short-term culture (8 days) (Fig. 2B) and accumulated more population doublings in long-term culture (100 days) compared to BMSCs (Fig. 2C). Both iMSCs and BMSCs expressed MSC-associated cell surface antigens, including CD29, CD44, and CD90, while lacking the hematopoietic-associated marker CD45 (Fig. 2D). To assess the differentiation potential of cells, iMSCs and BMSCs were induced for adipogenesis, chondrogenesis, and osteogenesis. Adipogenic induction resulted in the formation of lipid droplets detected by Oil red O staining and quantification (Fig. 2E) and a significant increase in the expression of the adipocyte-associated marker *LPL* (Fig. 2F), indicating adipocyte differentiation from both cell types. For chondrogenesis, iMSCs and BMSCs demonstrated the ability to form chondrocytes, as evidenced by Alcian blue staining for glycosaminoglycan (GAG) production (Fig. 2G). Quantification using the dimethylmethylene blue assay, normalized to DNA content, confirmed the presence of GAG in the induced cells.

**Figure 2.**
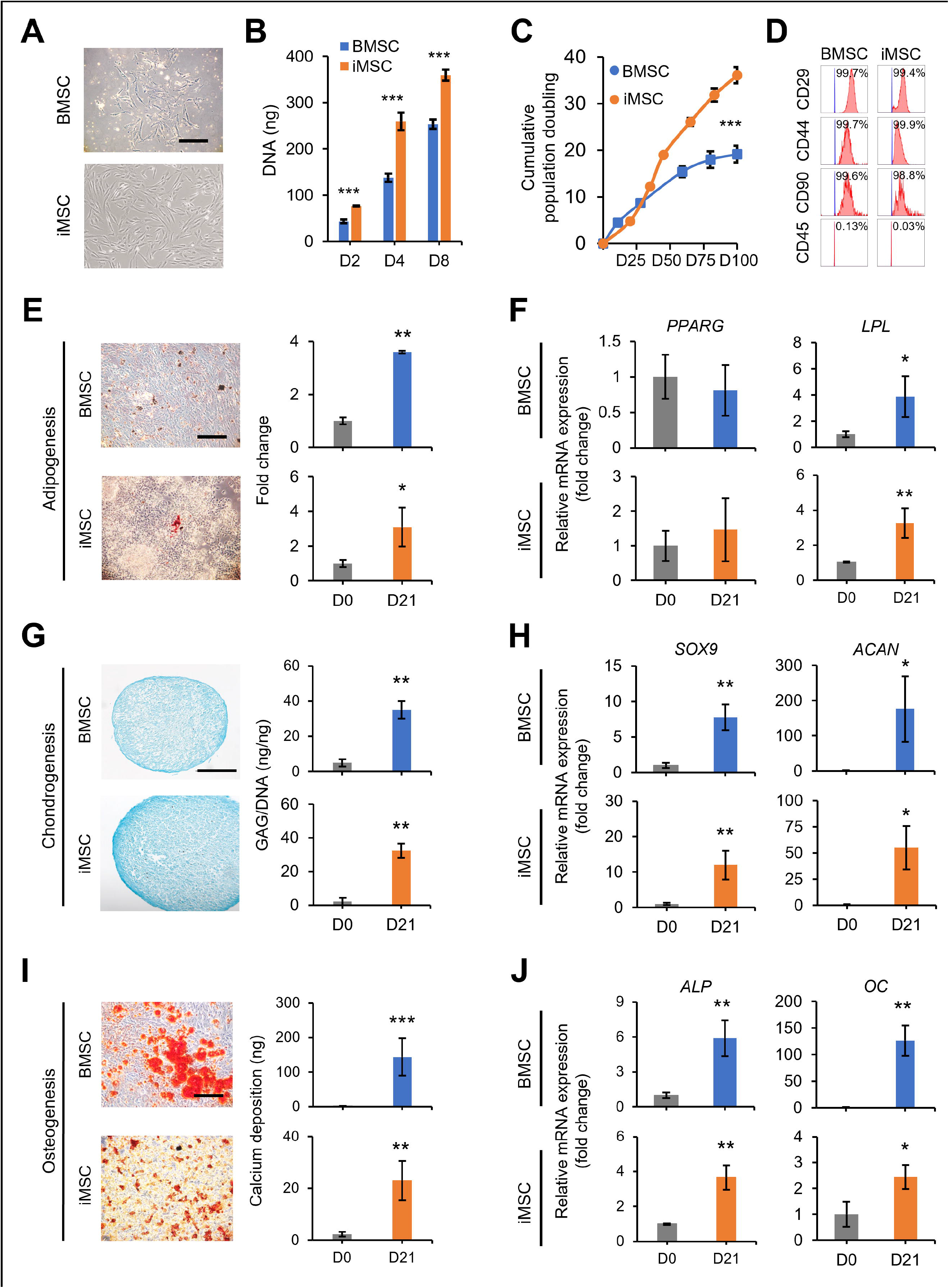
Characterization of minipig iMSCs and BMSCs. (A) Morphology of iMSCs and BMSCs. (B) Proliferation of iMSCs and BMSCs measured by DNA content at different time points. (C) Cumulative population doubling levels of iMSCs and BMSCs measured at each passage up to 100 days. (D) Flow cytometry analysis of iMSCs and BMSCs for detection of MSC-associated surface markers (CD29, CD44, and CD90) and hematopoietic marker (CD45). (E) Oil red O staining and quantification of lipid droplets following adipogenesis, and (F) transcript levels of adipose-associated markers (*PPARG* and *LPL*) in iMSCs and BMSCs after adipogenesis. (G) Alcian blue staining and quantification of glycosaminoglycans (GAGs) following chondrogenesis, and (H) transcript levels of cartilage-associated markers (*SOX9* and *ACAN*) in iMSCs and BMSCs after chondrogenesis. (I) Alizarin red S staining and quantification of calcium deposition following osteogenesis, and (J) transcript levels of bone-associated markers (*ALP* and *OC*) in iMSCs and BMSCs after osteogenesis. Scale bar = 200 µm. \**p* < 0.05, \*\**p* < 0.01, \*\*\**p* < 0.001; n = 3.

Moreover, chondrogenic induction led to increased expression of chondrocyte-specific markers *SOX9* and *ACAN* in both cell types (Fig. 2H). In terms of osteogenesis, both iMSCs and BMSCs exhibited the differentiation into osteoblasts, as indicated by Alizarin red staining for mineral deposition and calcium quantification (Fig. 2I). Additionally, the expression of osteocyte-associated markers *ALP* and *OC* was observed in both cell types upon osteogenic induction (Fig. 2J). These results demonstrated the multilineage differentiation potential of both iMSCs and BMSCs.

### *In vitro* induction of chondrogenesis in iMSC- and BMSC-laden fibrin glue/nanofiber constructs for generation of engineered cartilage implants

We manufactured a biomaterial construct consisting of fibrin glue hydrogel mixed with iMSCs or BMSCs enclosed by nanofibrous mats (Fig. 3A) and induced for chondrogenesis to generate an engineered cartilage implant for chondral defect repair. Live/dead assay demonstrated high cell viability in both iMSC and BMSC constructs with live cells stained green and dead cells stained red (Fig. 3B). Proliferation analysis based on total DNA content revealed distinct growth patterns in iMSC and BMSC constructs during chondrogenic induction. iMSC constructs exhibited an increase in cell number within the initial 7 days, while BMSC constructs remained stable (Fig. 3C). Subsequently, both iMSC and BMSC constructs underwent transformation into engineered cartilage-like implants after 21 days, characterized by a notable presence of GAG content (Fig. 3D). Importantly, the expression levels of key cartilage-associated markers, such as *SOX9*, *COL2A1*, and *ACAN*, were significantly upregulated in chondrocytes derived from both iMSC and BMSC constructs (Fig. 3E), affirming the successful generation of engineered cartilage implants in our in vitro model.

**Figure 3.**
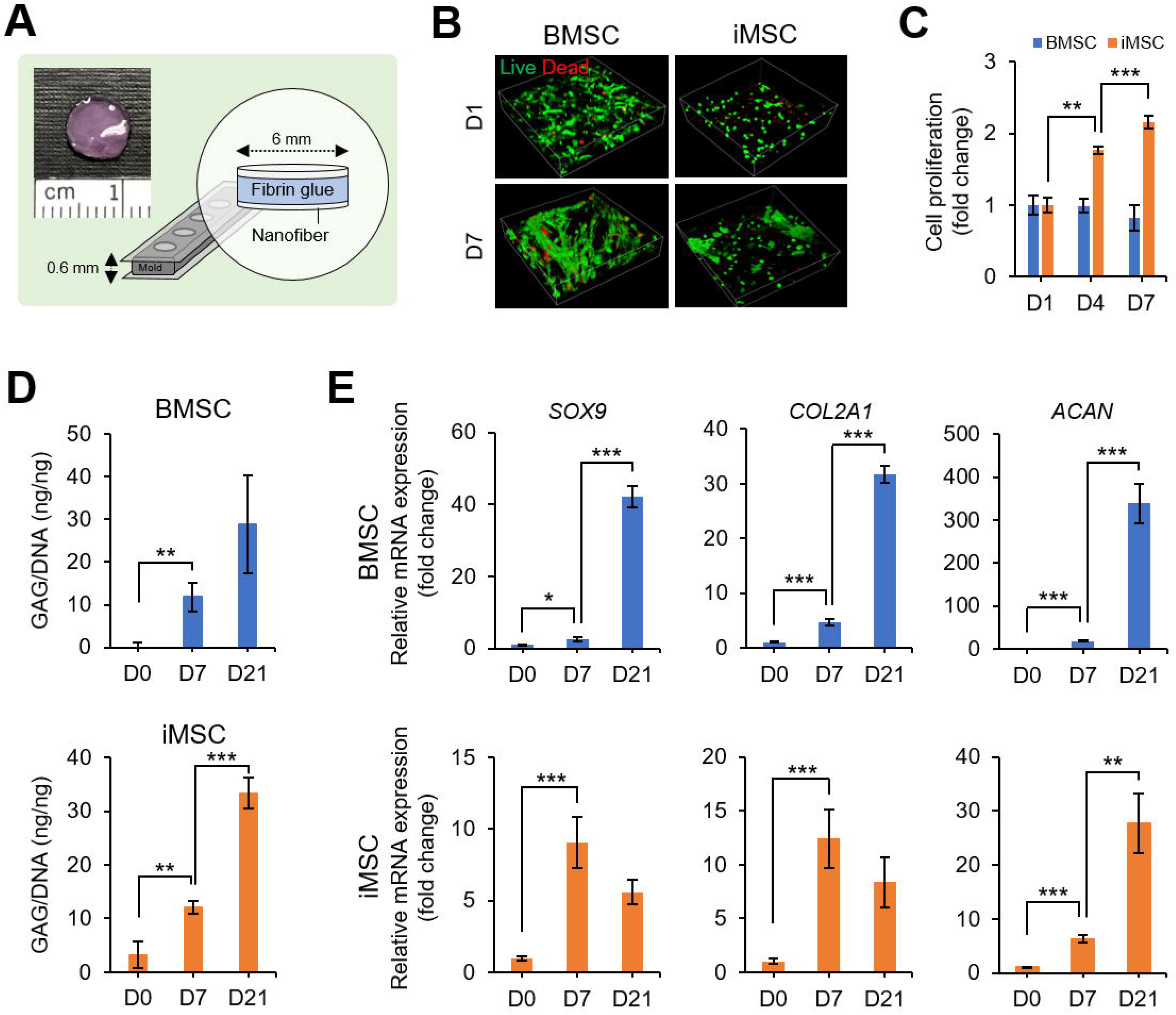
Chondrogenic evaluation of iMSCs and BMSCs seeded in fibrin glue/nanofiber constructs. (A) Schematic of the fibrin glue/nanofiber sandwich construct, where iMSCs or BMSCs are mixed with fibrin glue and assembled with nanofibrous mats to assess chondrogenic capacity. (B) Live (green) and dead (red) staining of iMSCs or BMSCs cultured in the fibrin glue/nanofiber construct at different time points. (C) Proliferation of iMSCs and BMSCs cultured in the construct, measured by DNA content at days 1, 4, and 7. (D) Quantification of GAG production per DNA content of iMSCs and BMSCs following chondrogenesis in the sandwich construct, measured at different time points. (E) Dynamics of the transcript levels of cartilage-associated markers (*SOX9*, *COL2A1*, and *ACAN*) during chondrogenic differentiation of iMSCs or BMSCs into chondrocytes at different time points. \**p* < 0.05, \*\**p* < 0.01, \*\*\**p* < 0.001; n = 3.

### Secure retention of engineered cartilage implants in chondral defects of minipig femoral condyles

To create a standardized chondral defect suitable for implanting engineered cartilage, we employed a specially designed hand-held trocar tool (Fig. 4A). This tool allowed us to create a circular lesion with a consistent depth, measuring 7 mm in diameter and 0.6 mm in depth, specifically on the weight-bearing surface of the medial femoral condyle. Importantly, we ensured that the subchondral bone remained intact and unaffected (Fig. 4B). Following the creation of the chondral defect, we introduced the engineered cartilage implant derived from either a BMSC- or iMSC-laden construct, or an acellular fibrin glue/nanofibrous mat construct. Prior to implantation, the cartilage implant was coated with fibrin glue to enhance its stability within the defect. For comparison, we included a control group that underwent the well-established microfracture technique, producing evenly spaced subchondral holes with a diameter of 1 mm (Fig. 4C). Postoperative MRI images taken after one month confirmed the secure placement of the implants within the targeted cartilage defect sites in the minipig femoral condyles (Fig. 4D).

**Figure 4.**
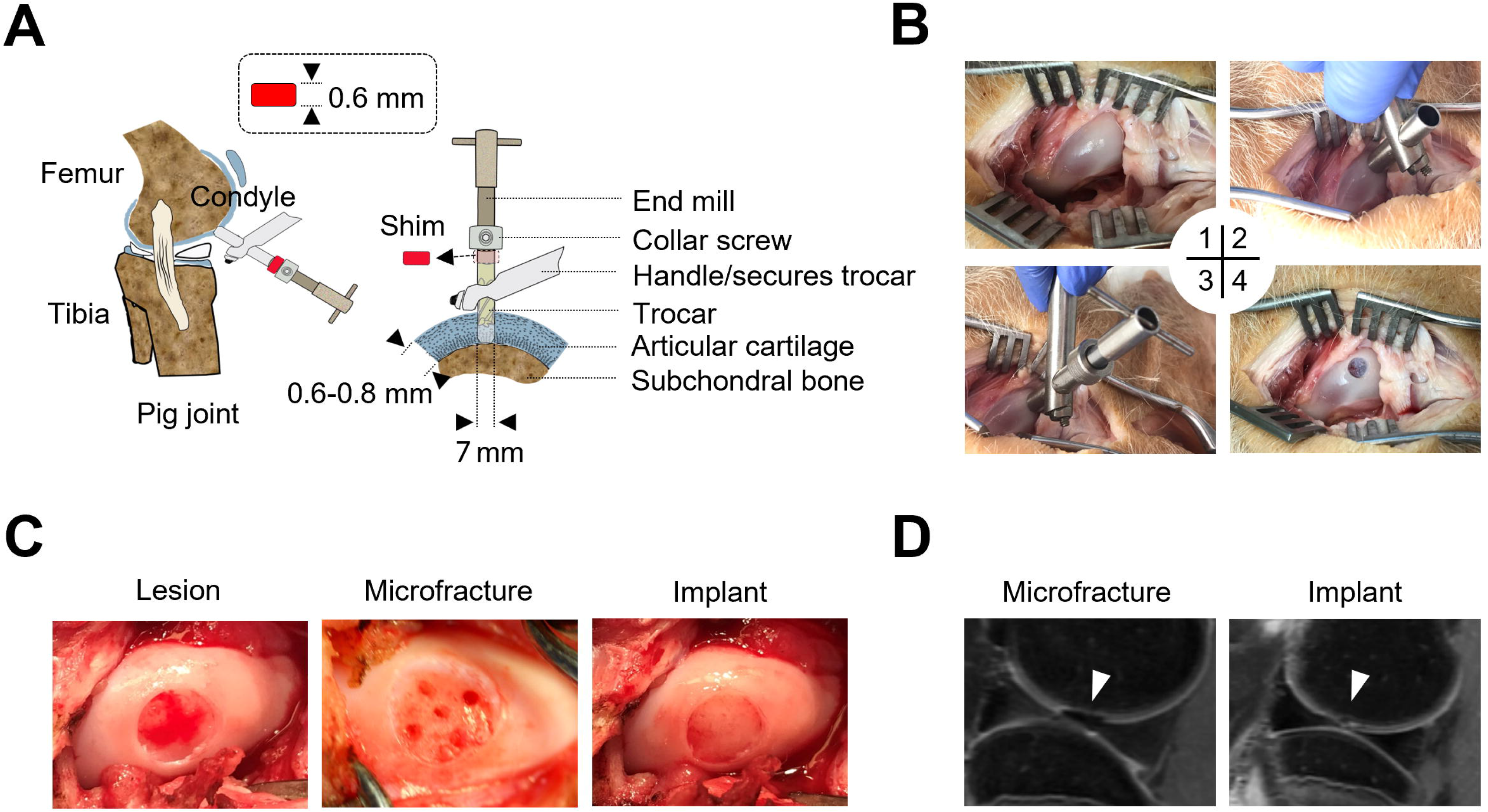
Chondral defects on medial femoral condyles and engineered cartilage construct implantation. (A) Illustration of the custom-designed trocar tool utilized to create a critical-sized articular cartilage defect, measuring 7 mm in diameter and 0.6 mm in depth, while preserving the integrity of the subchondral bone. (B) Step-by-step procedure for creating an articular cartilage defect in a minipig as follows: 1) A 10-cm skin incision was made to expose the defect site on the weight-bearing surface of the medial femoral condyle. 2) The trocar tool was placed on the surface of the articular cartilage. 3) The end mill was inserted into the trocar along with a 0.6 mm thick shim and a secure collar. 4) The shim was then removed, and gentle back and forth movements of the end mill were applied to create a critical-sized articular cartilage defect with a diameter of 7 mm and a depth of 0.6 mm. (C) Representative images of chondral lesions generated by the trocar tool prior to scaffold implantation or microfracture drilling (all groups; left), after microfracture drilling (microfracture group; middle), and after engineered construct implantation (acellular and cellular groups; right). (D) Magnetic resonance imaging (MRI) images captured 1 month after surgery demonstrating the repaired status of the defects in the microfracture and construct implantation groups.

### iMSC-Ch implants facilitate regeneration of hyaline cartilage in chondral defects of femoral condyles

Four months after the implantation surgery, we conducted a histological evaluation of critical-size chondral defects repaired by engineered cartilage. Macroscopic analysis of the harvested joints revealed partial filling of the cartilage defects in the cell-laden construct and microfracture groups, whereas the acellular construct group showed little to none neo-tissue generated in the defects (Fig. 5A). Quantitatively, the acellular group received significantly lower scores based on visual assessment using the ICRS-I scoring system compared to the other three groups (Fig. 5B).

**Figure 5.**
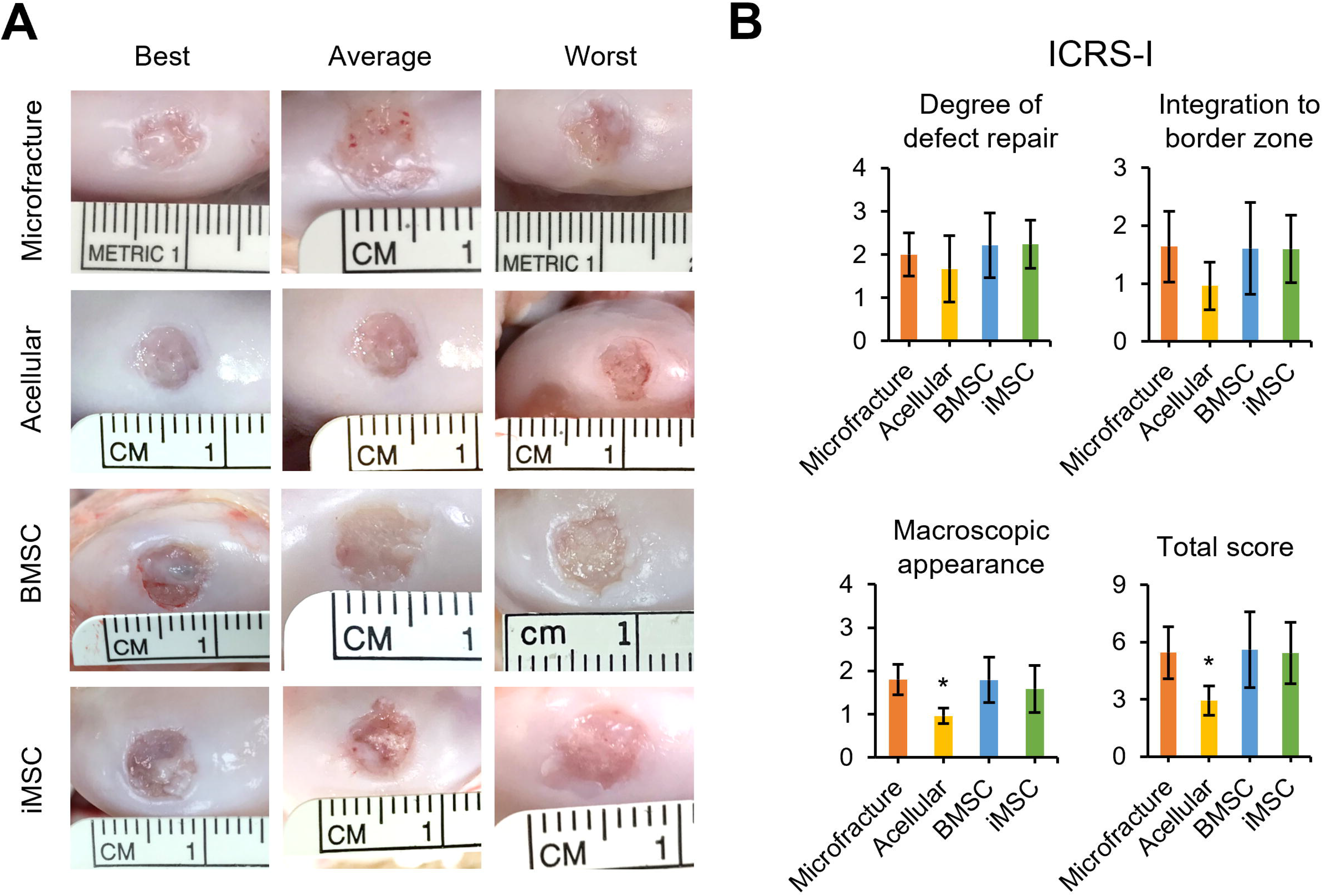
Macroscopic assessment of articular cartilage repair. (A) Representative macroscopic images of the repaired articular cartilage lesions in minipigs, taken 4 months after the surgical procedure. The images portray the repair outcomes of different treatment groups, including the microfracture group (clinical treatment control) and the acellular group (cellular treatment control). (B) The ICRS-I scoring system utilized for the visual assessment of the repaired articular cartilage in minipigs. The graph presents the scores obtained from the evaluation, with statistical significance indicated by asterisks (\**p* < 0.05). The data reflects a sample size of n = 6.

Histological analysis by hematoxylin and eosin (H&E) staining confirmed the generation of new tissue, partially covering the chondral defects in the BMSC-Ch, iMSC-Ch, and microfracture groups, while the acellular group showed minimal tissue repair (Fig. 6A).

**Figure 6.**
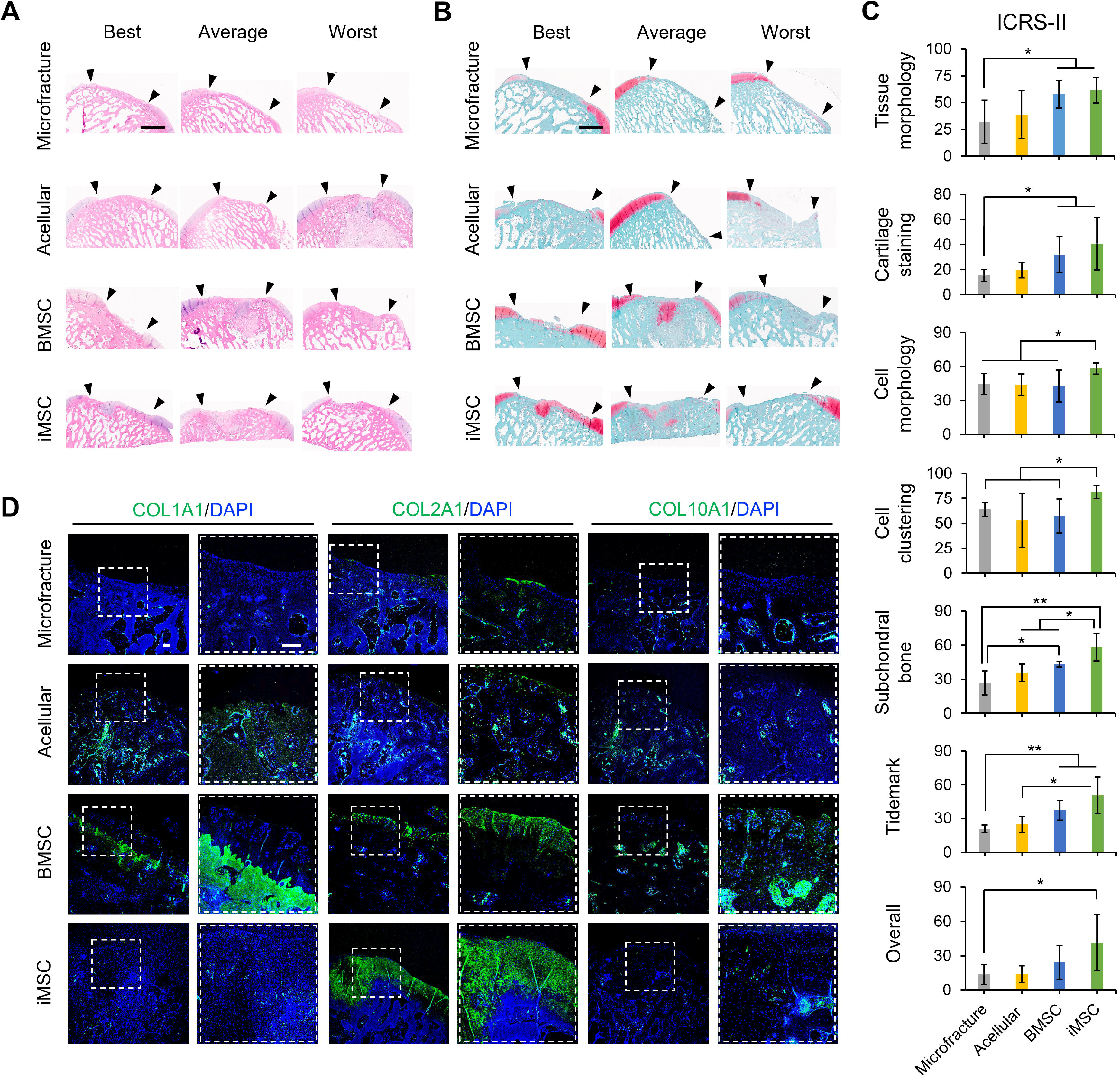
Histological assessment of articular cartilage repair. (A) H&E-stained images illustrating the histological assessment of the articular cartilage defect in minipigs, taken 4 months after the surgical procedure. The site of the cartilage defect creation is indicated between the black arrows. (B) Safranin O/Fast Green staining performed on minipig articular cartilage to analyze cartilage regeneration. In this staining, cartilage is displayed as red, while other connective tissues are stained green. The site of the cartilage defect creation is indicated between the black arrows. (C) The histological assessment of the regenerated minipig articular cartilage conducted using the ICRS-II grading score. The graph presents the scores obtained from the evaluation, with statistical significance indicated by asterisks (\**p* < 0.05, \*\**p* < 0.01). The assessment was performed on a sample size of n = 6. (D) Immunofluorescence staining performed on the regenerated minipig articular cartilage to detect the presence of cartilage-associated markers, including COL1A1, COL2A1, and COL10A1. The magnified images on the right represent regions enclosed by dashed boxes in the left columns. The nuclear DNA is labeled with DAPI. The scale bar in the images corresponds to a length of 200 µm. The data presented was obtained from a sample size of n = 6.

Alterations of the subchondral bone structure, likely caused by changes in the mechanical environment of a repaired joint, were found in some animals of each group. In terms of neo-cartilage matrix formation, Safranin O/Fast Green staining demonstrated more regenerated cartilage in the iMSC-Ch group compared to that in the BMSC-Ch group, whereas the microfracture and acellular groups showed little or no cartilage regeneration (Fig. 6B).

Quantitative histological assessment using the ICRS-II scoring system revealed that both the BMSC-Ch and iMSC-Ch groups achieved significantly higher scores in the parameters of tissue morphology, cartilage staining, subchondral bone, and tidemark compared to the microfracture control group (Fig. 6C). When comparing the ICRS-II scores between the two cellular implant groups, the repaired cartilage by iMSC-Ch implants received significantly higher scores in the parameters of cell morphology, cell clustering, and subchondral bone regeneration compared to that by BMSC-Ch implants. Immunofluorescence staining of cartilage-associated markers demonstrated that while there was a lack of regenerated cartilage matrix in the microfracture and acellular groups, the cellular implant groups showed the production of collagenous matrix. The tissue regenerated by iMSC-Ch implants was rich in collagen type 2 and had nearly undetectable collagen types 1 and 10. On the other hand, the BMSC-Ch group showed a substantial amount of collagen type 1, some collagen type 2, and detectable collagen type 10 (Fig. 6D). Taken together, our findings demonstrate that the cellular implant groups showed improved cartilage repair compared to the microfracture or acellular groups. Notably, iMSC-Ch implants led to the regeneration of hyaline cartilage-like tissue, while BMSC-Ch implants the formation of fibrocartilage-like tissue.

### Reduction of inflammation in joints repaired by iMSC-Ch implants

We conducted an analysis of inflammation-associated cytokines in the joint microenvironment of minipigs that received acellular or cellular implants. The microfracture group was excluded from the assay to ensure a suitable comparison among the implant groups that involved scaffolding biomaterial. Our findings revealed that BMSC-Ch implants significantly upregulated IFNG, ILRA, IL6, and IL12 in the synovial fluid of minipigs compared to acellular implants (Fig. 7A). When comparing the cellular implant groups, higher levels of proinflammatory factors, including IFNG, IL8, IL12, IL18, and TNFA, were observed in the synovial fluid of minipigs receiving BMSC-Ch implants compared to those receiving iMSC-Ch implants. Furthermore, the iMSC-Ch implant group demonstrated higher levels of anti-inflammatory factors, namely IL4 and IL1RA, compared to the BMSC-Ch implant group.

**Figure 7.**
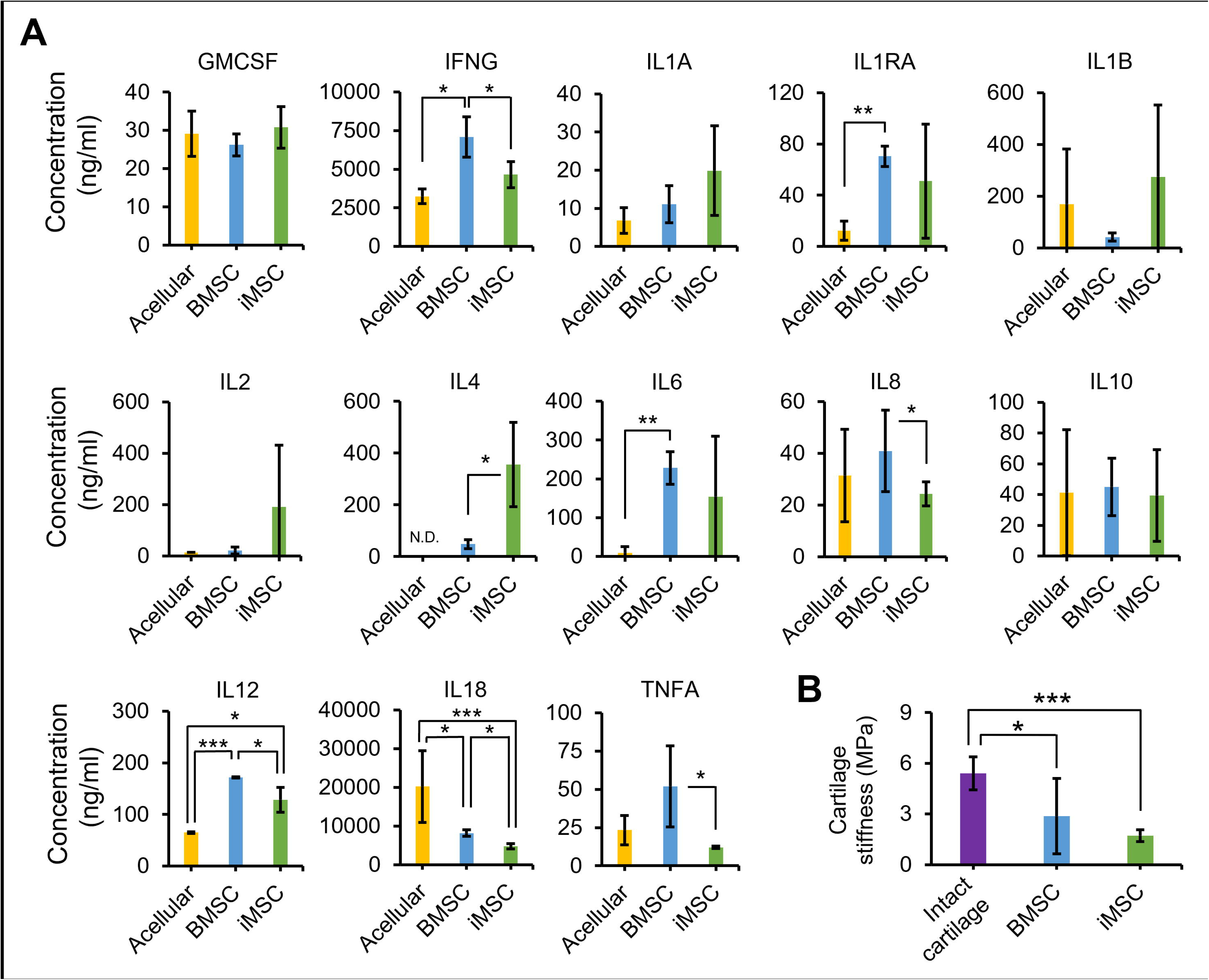
Inflammatory cytokine levels in synovial fluid and stiffness of regenerated cartilage in repaired joints. (A) Assessment of anti- and pro-inflammatory marker expression in synovial fluid from repaired joints of minipigs. The expression levels of these markers were evaluated to analyze the microenvironment within the repaired joints. Statistical significance is indicated by asterisks (\**p* < 0.05, \*\**p* < 0.01, \*\*\**p* < 0.001). The analysis was conducted on a sample size of n = 6. (B) The indentation hardness of both intact and repaired cartilage measured to assess their mechanical characteristics. Statistical significance is indicated by asterisks (\**p* < 0.05, \*\**p* < 0.01, \*\*\**p* < 0.001). The analysis was performed on a sample size of n = 6.

In order to assess the mechanical properties of the repaired cartilage, we measured the stiffness of both regenerated and intact cartilage as a reference control (Fig. 7B). Our results indicated that the stiffness of regenerated articular cartilage derived from iMSC-Ch implants was comparable to that of BMSC-Ch implants. However, it is worth noting that the regenerated cartilage exhibited significantly lower stiffness compared to intact cartilage. These findings suggest that iMSC-Ch implants possess the ability to modulate the inflammatory response in a repaired joint, although further enhancements in their mechanical properties are necessary.

## Discussion

The significance of this study is to evaluate the feasibility of using *ex vivo*-generated autologous iPSC-derived chondrocytes for repairing critical-sized chondral defects in a large animal model. To the best of our knowledge, this study is the first to address the question of whether the autologous iPSC strategy is viable for cartilage repair. We generated transgene-free minipig iPSCs using the nonintegrating episomal approach^24^ to demonstrate their potential for clinical applications. Additionally, we included groups of BMSCs and microfracture, representing the gold standard cell type and surgical intervention, respectively, as controls to compare the outcome of cartilage repair by iPSC derivatives. Our major findings revealed that iPSC-derived chondrocytes exhibit higher proliferation rates than BMSC-derived chondrocytes, while both types show a similar capacity for cartilaginous matrix production. Notably, autologous iMSC-Ch-laden implants promote hyaline cartilage formation and enhance the immune microenvironment of the joint compared to autologous BMSC-laden implants. These findings demonstrate the potential of autologous iPSCs as a source of therapeutic cells for articular cartilage repair.

The optimal choice between autologous and allogeneic cell sources for cartilage repair therapies remains a topic of ongoing investigation due to the distinct advantages and limitations associated with each approach. Autologous cell transplantation is considered a safer option compared to allogeneic transplantation. However, the high costs involved in generating patient-specific cells meeting cGMP requirements restrict the clinical feasibility of autologous approaches.^25^ On the other hand, allogeneic “off-the-shelf” graft implantation provides a more cost-effective alternative with readily available cell products and an abundant supply of cells.

Nevertheless, the risk of immune rejection presents a significant challenge to successful allogeneic cell transplantation,^26, 27^ even with HLA matching, necessitating the use of immunosuppressive medications that carry increased infection and cancer risks.^28^ Excitingly, a recent study conducted by Tsumaki’s group has demonstrated promising results in a primate animal model using allogenic iPSC-derived cartilage implants.^29^ These implants successfully filled chondral defects and transformed into white articular cartilage, resembling the surrounding tissue. These findings help address safety concerns associated with the use of allogenic iPSCs for cartilage reconstruction. However, it is essential to note that the previous study primarily focused on relatively small cartilage lesions measuring 1 mm in diameter and 0.5 mm in depth. Further investigation is necessary to determine if similar repair outcomes can be achieved in a large animal model with critical-sized defects, as employed in our study. Studying critical-sized defects is crucial as they present greater challenges in terms of mechanical load distribution and inflammation associated with the lesion, which necessitate achieving satisfactory repair outcomes.

In this study, we implemented autologous cell implants and transgene-free cellular reprogramming to address safety concerns associated with immunogenicity and tumorigenicity in iPSC-derived cell therapies.^30, 31^ Our findings indicate that minipigs receiving cartilage implants generated from their own BMSC- or iPSC-derived chondrocytes experienced minimal immune response-related inflammation and pain. Moreover, no tumor formation was observed at the defect site after iMSCs implantation. The utilization of an integration-free reprogramming method likely contributed to this positive outcome. A report suggests that integrating methods using recombinant DNA or viral vectors can lead to genomic disruptions and oncogenic activations.^32^ In contrast, non-integrating methods like episomal plasmids, although having lower transfection efficiencies, are considered reliable and safe for translational research and future clinical applications. It is important to note that the pMaster12 plasmid utilized in this study for iPSC reprogramming contains the MYC transcription factor. MYC can enhance cell reprogramming efficiency by regulating cell metabolism during early reprogramming stages through the induction of a robust hybrid energetics program,^33^ although it has been shown to be associated with the promotion of undesired tumor formation. Our study confirmed that all iMSC lines were transgene-free, indicating the absence of the four exogenous Yamanaka factors. This outcome alleviates safety concerns regarding the presence of these factors in the implanted cells.

Another noteworthy aspect of this study is the utilization of a skeletally mature translational model for chondral defect repair. It has been observed in previous studies that articular cartilage in younger models has a higher capacity for self-regeneration compared to older models.^34, 35^ For example, Varasa et al. reported that articular cartilage of skeletally immature pigs retains the ability for spontaneous repair.^36^ Their study showed no significant difference in cartilage repair between the autologous chondrocyte implantation group and the sham group in pigs treated at 8 months of age, as evaluated at 3 and 12 months follow-ups. To mitigate the potential confounding effect of spontaneous cartilage repair, which is often observed in immature minipigs, we allowed the animals to reach 24 months of age before performing the experimental surgery. It is noteworthy that the acellular control group demonstrated the least favorable outcomes in terms of cartilage repair, aligning with the existing understanding that adult cartilage possesses limited regenerative potential. By employing skeletally mature animals in our experimental design, we aimed to provide a more objective evaluation of cartilage repair that closely resembles the clinical scenario in adult human patients.

The current findings are in line with our previous research, demonstrating the superior cartilage repair capacity of human iMSCs compared to their parental MSCs derived from synovial fluid in a spontaneous osteoarthritis model.^37^ However, conflicting results from other studies suggest that BMSCs may have higher chondrogenic potential than iPSC derivatives. For example, Diederichs et al. compared iPSC-derived MSC-like progenitor cells (iMPCs) with parental BMSCs and found that iMPCs showed higher expression of cartilage-associated markers and greater GAG production after chondrogenesis induction.^38^ The disparity in results could be due to differences in the induction procedures for MSCs. These variations may lead to the induction of distinct MSC subpopulations with different chondrogenic differentiation capacities. In contrast, our study demonstrated that iMSC-Chs not only exhibited higher chondrogenic potential in repairing cartilage lesions but also generated hyaline-like cartilage, as evidenced by predominant expression of the hyaline cartilage-associated marker COL2A1 and minimal expression of the hypertrophic chondrocyte-associated marker COL10A1. These findings align with a recent study by Pothiawala et al., which identified a subpopulation of iPSC derivatives capable of generating and maintaining a “true” permanent-like articular cartilage phenotype that does not progress towards endochondral ossification.^39^ They identified a chondroprogenitor population positive for GDF5 derived from iPSC-derived mesenchymal cells along the paraxial mesodermal lineage. Collectively, our study highlights the potential of iPSC derivatives in generating hyaline-like cartilage for the repair of joint defects.

While the potential of iPSC derivatives for cartilage regeneration has been extensively investigated, including in our study, there remains a common limitation of small sample sizes among reported studies. This limitation contributes to variable results observed among animals. Additionally, it is worth noting that MSCs may not be the optimal cell type for repairing chondral defects due to their inherent propensity towards endochondral ossification. Recent studies have shown promising outcomes by utilizing chondrocytes derived directly from iPSCs, bypassing the MSC stage.^40, 41^ This approach potentially avoids the hypertrophy often observed in MSC-derived chondrocytes and may offer a more direct and effective method for generating hyaline cartilage. However, further investigation and comparison between these two strategies are necessary to determine the most suitable approach for successful cartilage regeneration.

In conclusion, this study demonstrates the potential of autologous iPSCs as a viable source of therapeutic cells for repairing articular cartilage defects. We provide evidence that iPSC-derived chondrocytes show comparable cartilaginous matrix production to BMSC-derived chondrocytes while exhibiting higher proliferation rates. Moreover, autologous iMSC-Ch-laden implants promote hyaline cartilage formation and enhance the immune microenvironment of the joint. These findings highlight the promising role of iPSC derivatives in cartilage repair and their potential as a safe and effective therapeutic strategy. Future studies should aim to optimize the chondrogenic potential of iPSC derivatives and investigate their long-term effects in larger animal models to further validate their clinical translatability.

## Materials and Methods

### Isolation of minipig fibroblasts and BMSCs

The animals used in this study and the experimental procedures were approved by the University of Wisconsin-Madison Institutional Animal Care and Use Committee. To generate autologous iPSCs, fibroblasts for cellular reprogramming were isolated from ear notch samples taken from 6 Yucatan minipigs at the age of 3 months, following our previously published protocol.^24^ In brief, the collected dermal tissue was digested in buffer medium supplemented with collagenase/dispase (MillporeSigma, Burlington, MA) for 2 hours. After digestion, growth culture medium composed of low-glucose Dulbecco’s Modified Eagle Medium (DMEM; Thermo Fisher Scientific, Waltham, MA, USA), 10% fetal bovine serum (FBS; Atlanta Biologicals, Atlanta, GA, USA), and 1% antibiotics (Thermo Fisher Scientific) was added to terminate the digestion process. The solution was then filtered through a 70-µm Falcon Cell Strainer (Corning, Glendale, AZ, USA) to remove undigested tissue, and cells were collected by centrifugation at 300 x g for 5 minutes. The cells were resuspended in growth culture medium and plated in a T75 culture flask, and then maintained in a humidified atmosphere of 5% CO_2_ and 95% air at 37°C. When the cells reached 70-80% confluence, they were trypsinized using 0.05% trypsin/EDTA (Thermo Fisher Scientific) and replated at a density of 1000 cells/cm^2^. The growth culture medium was changed every 3 days.

To harvest bone marrow from the iliac crest for BMSC isolation, 5 Yucatan minipigs were anesthetized using a combination of Telazol (5 mg/kg), ketamine (5 mg/kg), and glycopyrrolate (0.01 mg/kg) administered intramuscularly (IM). Propofol was given intravenously to the animals for intubation, followed by 2-3% isoflurane for the maintenance of general anesthesia. Preoperative doses of the antibiotic procaine penicillin G (40,000 IU/kg) and the pain medication buprenorphine (0.02 mg/kg) were administered IM. The minipigs were then transferred to a surgical suite and positioned in lateral recumbency. A 5-cm skin incision was made over a randomly selected iliac crest to expose the iliac bone using periosteal elevators. A 2− mm diameter hole was drilled on the dorsal portion of the iliac wing. Using a heparin (100 IU/ml) coated 10-ml syringe attached to a 2-mm diameter biopsy needle, 2-3 ml of bone marrow fluid was collected from the iliac bone marrow cavity. The layers of muscle, fascia, and sub-skin were closed with an absorbable suture (#4 Polysorb), and the skin was closed with a non-absorbable suture (#3 Surgipro). A second dose of procaine penicillin G (40,000 IU/kg) was administered immediately postoperatively, prior to recovery. The isolated bone marrow fluid was transferred into a T75 flask and cultured with low glucose DMEM growth medium containing 10% FBS and 1% antibiotics, and maintained at 37°C in a humidified atmosphere of 5% CO_2_.

### Generation of minipig fibroblast-derived iPSCs and iMSCs

To generate minipig iPSCs, we followed the protocol of our previous study^20^ to transfected 6 individually isolated fibroblast lines with 4 μg of episomal plasmid (#58527; Addgene, Watertown, MA, USA) using the NucleofectorTM II with the A-024 program (Amaxa, Walkersville, MD, USA). The transfected cells were then seeded into a 6-well plate containing an irradiated mouse embryonic fibroblast (MEF; WiCell, Madison, WI, USA) feeder layer, with modified E8 medium (E8 medium (Thermo Fisher Scientific) supplemented with 5 ng/ml activin A (Thermo Fisher Scientific), 1.5 µM CHIR99021 (STEMCELL Technologies, Vancouver, BC, Canada), 2.5 µM IWR-1 (STEMCELL Technologies), and 10 ng/ml LIF (Thermo Fisher Scientific)), for 21 days to induce iPSC colonies. From Day 2 to Day 7, G418 (400 µg/ml) was added to the modified E8 medium to enhance reprogramming efficacy. Under a microscope, iPSC colonies were hand-picked using a 27G needle and individually transferred to fresh 24-well plates containing an MEF feeder layer and modified E8 medium for expansion.

The iPSCs were characterized using Fast Blue RR Salt (Sigma Aldrich, St. Louis, MO, USA) staining to detect alkaline phosphatase and flow cytometry to determine the expression of pluripotency markers. Furthermore, iPSCs were differentiated into ectodermal, endodermal, and mesodermal lineage cells *in vitro* using the STEMdiff™ Trilineage Differentiation kit (STEMCELL Technologies) according to the manufacturer’s instructions. Subsequently, qRT-PCR was performed to analyze the mRNA expression of representative iPSC markers and germ layer lineage-specific identifiers using the primers listed in Table 1.

**Table 1.**
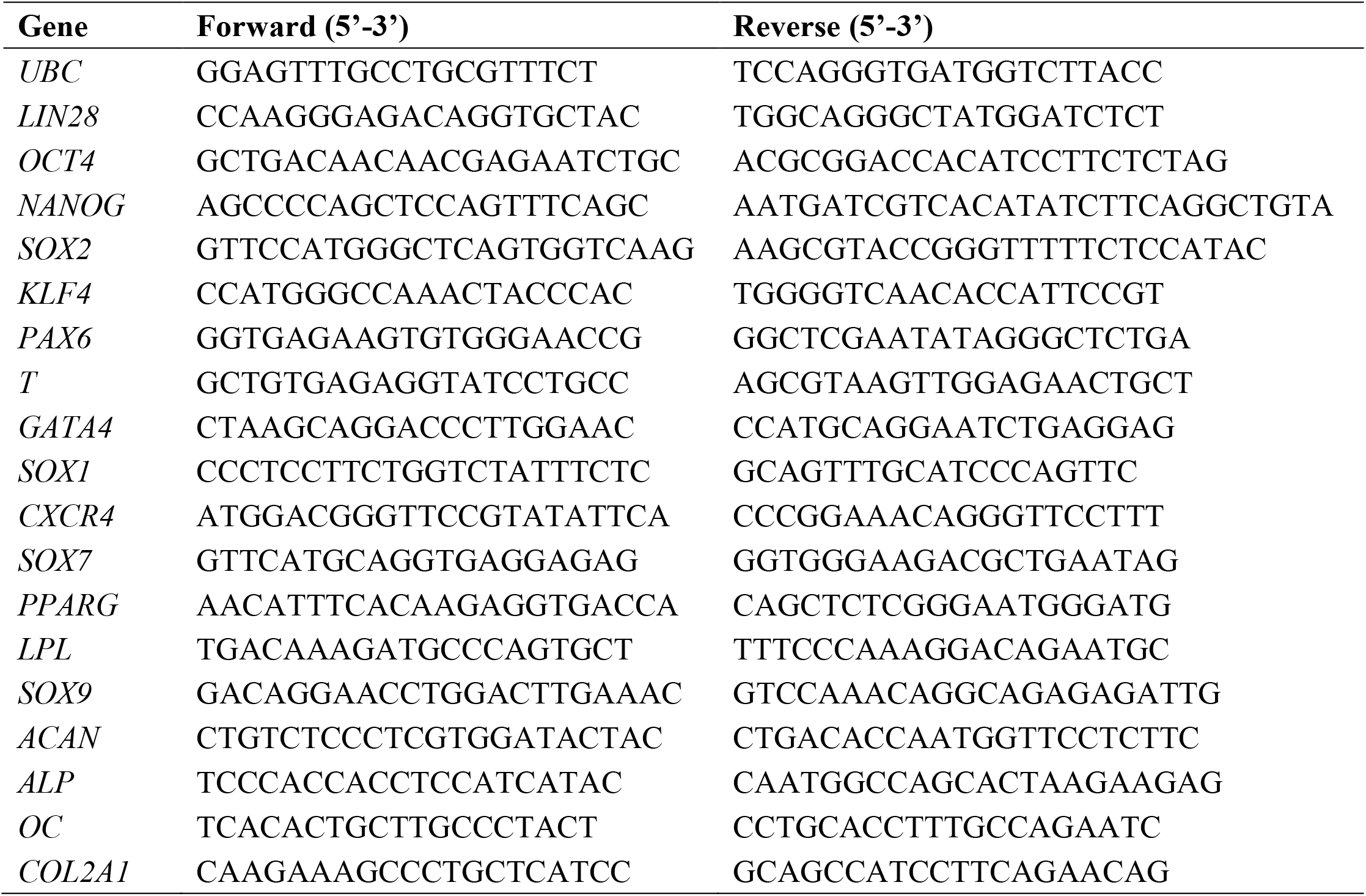
Minipig Primer Sequences for Quantitative PCR Analysis.

All six independent iPSC lines were further differentiated into iMSCs using the STEMdiff TM Mesenchymal Progenitor kit (STEMCELL Technologies) following the manufacturer’s instructions. Briefly, iPSCs were cultured in 24-well plates containing an MEF feeder layer with E8 medium until they reached 70-80% confluence. The culture medium was switched to STEMdiff™-ACF Mesenchymal Induction Medium (STEMCELL Technologies) for 4 days to induce iMSCs before the medium was changed to Complete MesenCult™-ACF Plus Medium (STEMCELL Technologies) for an additional 2 days and then the cells were transferred into a new 24-well plate without an MEF feeder layer. The cells underwent another 4 passages using the Complete MesenCult™-ACF Plus Medium. Finally, the culture medium was switched to low glucose DMEM growth medium containing 10% FBS and 1% antibiotics to maintain iMSCs. Daily media change was performed throughout the process.

### Cell proliferation

Proliferation of iMSCs and BMSCs seeded with or without fibrin glue/nanofiber constructs was assessed on Days 1, 4, and 7 in culture. DNA extraction from the cells was performed using Proteinase K (Millipore Sigma, Burlington, MA, USA), and the DNA content was quantified using the PicoGreen assay (Thermo Fisher Scientific), following the manufacturer’s instructions. Cumulative population doubling analysis was performed to evaluate the growth rate of long-term cell culture. The iMSCs and BMSCs were cultured and passaged when they reached 70-80% confluency for a duration of up to 100 days. The cell number at each passage was recorded and calculated. The cumulative number of population doublings was obtained by adding the calculated number from each passage to the number from the previous passage.

### Flow cytometric analysis

To characterize iPSC and MSC markers, cells were trypsinized, suspended, and washed three times using ice-cold washing buffer. The washing buffer contained 1% bovine serum albumin (Millipore Sigma), 5 mM EDTA (Thermo Fisher Scientific), and 25 mM HEPES (Thermo Fisher Scientific) in D-PBS (Cytiva, Marlborough, MA, USA). For iPSCs, the cells were incubated with primary antibodies (Thermo Fisher Scientific), including rabbit anti-OCT4, rabbit anti-NANOG, and rabbit anti-SOX2, for 30 minutes at 4°C. Subsequently, the cells were washed three times with the washing buffer and then incubated with the secondary antibody (Thermo Fisher Scientific), Alexa Fluor® 555 donkey anti-rabbit IgG, for an additional 30 minutes at 4°C. On the other hand, iMSCs and BMSCs were incubated with Alexa Fluor® 647 mouse anti-CD29 (Thermo Fisher Scientific), Alexa Fluor® 488 mouse anti-CD44 (Thermo Fisher Scientific), FITC mouse anti-CD90 (BD, Franklin Lakes, NJ, USA), and PERCP-CYTM5.5 mouse anti-CD45 (BD) for 30 minutes at 4°C. After three consecutive washes, the fluorescent cells were analyzed using Attune NxT Flow Cytometer (Thermo Fisher Scientific), and the data of marker expression were analyzed using FlowJo software (Tree Star, Ashland, OR, USA) following the manufacturer’s instructions.

### Induction of multilineage differentiation of iMSCs and BMSCs in culture

iMSCs and BMSCs were induced to differentiate into the adipogenic, chondrogenic, and osteogenic lineages for 21 days following our previously published protocol.^42^ For adipogenesis, cells were plated in a 6-well plate at a density of 100,000 cells/cm² and induced by adipogenic medium consisting of high-glucose DMEM, 10% FBS, 1% antibiotics, 1 µM dexamethasone (Millipore Sigma), 0.5 mM 3-isobutyl-1-methylxanthine (Millipore Sigma), and 1 µg/ml insulin (Millipore Sigma). For chondrogenesis, 5 x 10⁵ iMSCs or BMSCs were collected and centrifuged at 600 x g for 5 minutes to form cell pellets. The pellets were then induced by chondrogenic medium consisting of high-glucose DMEM supplemented with 1% antibiotics, 1% ITS + Premix (Corning), 0.9 mM sodium pyruvate (Sigma Aldrich), L-ascorbic acid-2-phosphate (50 g/ml, Sigma Aldrich), L-proline (Sigma Aldrich), 0.1 M dexamethasone (Sigma Aldrich), and TGFB1 (10 ng/ml) (Thermo Fisher Scientific). For osteogenesis, cells were seeded in a 6-well plate at a density of 5 x 10³ cells/cm² and cultured with osteogenic medium containing low-glucose DMEM, 10% FBS, 1% antibiotics, 10 mM β-glycerophosphate (Sigma Aldrich), 50 μg/ml l-ascorbic acid-2-phosphate, and 0.1 µM dexamethasone. During the differentiation period, the induction medium was changed every 3 days.

After 21 days of differentiation, the cells were analyzed for mRNA expression of fat-, cartilage-, and bone-associated markers by qRT-PCR with primers listed in Table 1, histological staining, and biochemical assays. Cells induced for adipogenesis were fixed with 10% neutral buffered formalin and stained by Oil Red O (Millipore Sigma) to detect lipid droplets. The staining was imaged and then quantified by dissolving it in 100% isopropanol for the absorbance measurement at a wavelength of 515 nm to determine the lipid content. Cell pellets induced for chondrogenesis were fixed with 4% formaldehyde, dehydrated, infiltrated with xylene, and embedded in paraffin. The paraffin blocks were sectioned into 8-μm sections, deparaffinized, and stained with Alcian blue (Polysciences, Warrington, PA, USA) to detect proteoglycans. The quantification of GAG was performed by digesting the cell pellets in papain solution and detected by the dimethylmethylene blue assay. The total GAG amount was normalized to the DNA content determined separately using the PicoGreen assay. For the analysis of osteogenesis, the cells were fixed with 60% isopropanol and stained by Alizarin Red S (Rowley Biochemical, Danvers, MA, USA) to detect calcium deposition. The calcium content was quantified by mixing the cells with 0.5 M hydrochloric acid (Sigma Aldrich) and measuring the mixture using the LiquiColor kit (Stanbio, Boerne, TX, USA) following the manufacturer’s instructions.

### Total RNA extraction and qRT-PCR analysis

For the analysis of mRNA expression of genes of interest, the Quick-RNATM MicroPrep kit (Zymo Research, Irvine, CA, USA) was used to extract total RNA from cells. The quantity and quality of the total RNA were then measured by the Infinite M Plex system (Tecan, Mannedorf, Switzerland) to prepare complementary DNA (cDNA) using the High-Capacity cDNA Reverse Transcription kit (Thermo Fisher Scientific). For real-time qPCR analysis, the cDNA samples were amplified with iQSYBR Green Premix (Bio-Rad, Hercules, CA, USA) and primers to detect specific genes of interest listed in Table 1. To determine the relative mRNA expression level of a target gene, the 2^−ΔCt^ method was employed and by which differences in cycle threshold (Ct) values between the target gene and the housekeep *GAPDH* were calculated.

### Fabrication of cell-laden fibrin glue/nanofiber constructs

To make biomaterial scaffolds for seeing cells as implants, electrospun ultrafine nanofibers were produced following our previously published protocol.^43^ Briefly, 15% (w/v) polymer solution was prepared by dissolving 90% poly-L-lactic acid (PLLA) (molecular weight = 50,000) (Polysciences, Warrington, PA, USA) and 10% polycaprolactone (PCL) (molecular weight = 70,000 - 90,000) (Sigma Aldrich) in hexafluoro-2-propanol (Fluka Analytical, St. Louis, MO, USA). The solution was loaded into a 10 ml syringe fitted with a 20-gauge needle and pumped out at a speed of 1 ml/hr during electrospinning with the setup of 12-kV DC and a 15-cm electrostatic field to produce nanofibers.

For the production of three-layered sandwich constructs, a custom-made stainless-steel mold set consisting of two flat rectangular plates and one perforated 6-hole plate (Fig. 3A) was employed. Each construct comprised a central component, a fibrin glue cylinder measuring 7 mm in diameter and 0.6 mm in height, surrounded by two 7-mm circular nanofibrous mats positioned above and below the fibrin glue cylinder. To create a cellular fibrin glue/nanofiber construct, iMSCs or BMSCs were trypsinized and suspended in thrombin (Baxter, Deerfield, IL, USA) at a concentration of 20 x 10^6^ cells/ml. In the mold, the bottom piece of the nanofibrous mat on a metal plate was infused with the thrombin-cell solution in a well. A combination of 5 μl of Sealer Protein Concentrate and 5 μl of the thrombin-cell solution was then added to the filled well before the top piece of the nanofibrous mat, infused with the thrombin-cell solution, was placed on top and covered with another metal plate. The resulting cellular constructs were subsequently transferred into 15-ml conical tubes (Corning) containing chondrogenic medium and underwent a 7-day induction period prior to surgical implantation. The medium was changed every 3 days during chondrogenic differentiation.

### A trocar tool designed to generate articular cartilage defects

A specialized trocar tool was custom-designed and constructed to facilitate the creation of an articular cartilage defect on the medial femoral condyle of a minipig. The trocar tool consisted of various components made from autoclavable stainless steel. To ensure stability during the procedure, a handle was incorporated into the design to securely position the trocar on the cartilage surface. The trocar tool featured an end mill, which was equipped with a stop collar and shim to regulate the depth of the defect. By employing controlled and gentle manual movements of the end mill, a precise and uniform articular cartilage defect was generated, maintaining the integrity of the underlying subchondral bone. The result was a smooth and flat surface at the base of the defect, ensuring consistency in the experimental procedure.

### Implantation of autologous iMSC- and BMSC-constructs for repair of chondral defects of minipig stifle condyles

The *in vivo* study was approved by the University of Wisconsin-Madison Institutional Animal Care and Use Committee. A total of nineteen 24-month-old Yucatan minipigs were randomly assigned to 4 subgroups: microfracture (n = 5), acellular (n = 3), autologous BMSC (n = 5), and autologous iMSC (n = 6). A combination of Telazol (6 mg/kg) and Xylazine (2.2 mg/kg) was administered to the animals as general anesthesia prior to surgical procedures, and the animals were then given isoflurane (1-3%) and 100% oxygen at a flow rate of 2 L/min through a semi-closed circuit during surgery. The minipigs were also given Excede® (5 mg/kg), a long-acting broad-spectrum antibiotic, and buprenorphine SR (0.24 mg/kg) to relieve pain. Once under anesthesia, the minipigs were positioned in dorsal recumbency and their hind limbs were suspended and prepared in a sterile manner through 3 rounds of saline solution rinsing followed by application of 4% hibiclens. A 5-cm skin incision was made to expose a medial femoral condyle. A specially designed custom-made trocar was used to create a cylindrical cartilage-only defect measuring exactly 7 mm in diameter and 0.6 mm in depth on the weight-bearing surface of the medial femoral condyle while preserving the subchondral bone of a minipig.

For the microfracture group, 5 equally spaced 1-mm diameter holes were drilled on a cartilage defect, penetrating the subchondral bone to disrupt blood vessels to allow clot formation. For the other 3 groups, acellular, BMSC-laden, and iMSC-laden constructs were implanted at created cartilage defects and fibrin glue (Baxter) was applied to securely hold the implants in place. The layers of muscle, fascia, and sub-skin were closed using #4-0 Polysorb suture material, and the skin was closed using #3-0 Surgipro suture material. One month after the surgery, imaging analysis by a 3.0 T MRI scanner (GE Healthcare Discovery MR750, Waukesha, WI, USA) was performed to examine the implants within the cartilage defects. The animals were kept for a period of 4 months before being euthanized for tissue harvesting and further analysis.

### Assessment of mechanical property of regenerated cartilage

The mechanical property of neo-cartilage generated at defects was determined by the measurement of tissue stiffness using the hand-held arthroscopic indentation probe Artscan 200 (Artscan Oy, Helsinki, Finland) in accordance with the manufacturer’s instructions. The Young’s modulus (E) of the tissue was calculated based on the equation E = F(1-ν)Rχ / (4α^3^κ), where Frepresents the force measured from the probe, ν is the Poisson’s ratio (assumed to be 0.5), R denotes the radius of curvature of the indenter (0.35 mm), α corresponds to the height of the indenter (0.13 mm), and κ represents the theoretical correction factor.^44^

**Analysis of cytokines in joint cavities and histological assessment of repaired cartilage** Synovial fluid and stifle joints of minipigs were collected 4 months posttreatment for the analysis of cytokines and histological evaluation, respectively. The collected synovial fluid was stored at −80°C before analysis. Levels of inflammation-associated cytokines were assessed by MILLIPLEX MAP Porcine Cytokine/Chemokine Magnetic Bead Panel – Immunology Multiplex Assay (Millipore Sigma), following the manufacturer’s instructions.

Stifle joint specimens were fixed in 10% neutral-buffered formalin and decalcified in 10% neutral-buffered EDTA. After the completion of decalcification was confirmed by radiography (Faxitron UltraFocus DXA, Tucson, AZ, USA), the specimens were embedded in paraffin and sectioned into 8-μm slices for subsequent immunohistochemical and histological staining. To stain samples, the sectioned slices were deparaffinized and subjected to unmasking by 0.1% (w/v) pepsin (Sigma Aldrich) in 0.01 N HCl for 15 minutes. Subsequently, permeabilization was performed using 0.1% Triton X-100 (Thermo Fisher Scientific) in D-PBS for 30 minutes, followed by blocking with ice-cold 0.1% (w/v) BSA in D-PBS for 30 minutes at room temperature. The specimen slides were then incubated with the primary antibodies, goat anti-human COL1A1 (Santa Cruz Biotechnology, Dallas, TX, USA), COL2A1 (Santa Cruz Biotechnology), and COL10A1 (Santa Cruz Biotechnology) at a 1:200 ratio for 1 hour at 4°C. Afterward, the slides were washed 3 times with ice-cold D-PBS to remove unbound antibodies and then incubated with the secondary antibody FITC donkey anti-goat (Santa Cruz Biotechnology) at a 1:200 ratio for 30 minutes at 4°C. Following 3 additional washes with ice-cold D-PBS to remove unbound secondary antibody, the slides were mounted with coverslips using mounting medium containing DAPI (Vector, Burlingame, CA, USA). Fluorescence imaging of the specimens was conducted using a Nikon A1R-s confocal microscope (Nikon, Tokyo, Japan). To quantitatively assess cartilage repair, optical macrographs and histological micrographs of the specimens were evaluated by 4 reviewers independently, who were blinded to the experimental groups. The ICRS-I and ICRS-II scoring systems were employed for the assessment of macrographs and micrographs, respectively. The reported scores in the results section represent the mean values calculated from assessments made by the independent reviewers.

## Statistical analysis

All the results presented in this study were derived from a minimum of three independent biological replicates (n = 3). Quantified data were expressed as means ± standard deviation (SD), and statistical comparisons between groups were performed using Student’s t-test or one-way analysis of variance (ANOVA) followed by a post hoc Tukey’s test. A significance level of *p* < 0.05 was considered statistically significant and used to determine the presence of significant differences among the experimental groups.

## Acknowledgements

This study was funded by the National Institute of Arthritis and Musculoskeletal and Skin Diseases of the National Institutes of Health (NIH) under Award Numbers R01 AR064803. The funder played no role in study design, data collection, analysis and interpretation of data, or the writing of this manuscript. The content is solely the responsibility of the authors and does not necessarily represent the official views of the National Institutes of Health.

